# Light regulated SIK1 remodels the synaptic phosphoproteome to induce sleep

**DOI:** 10.1101/2021.09.28.462159

**Authors:** Lewis Taylor, Teele Palumaa, Paul K Reardon, Steven Walsh, Bradley H Johnson, Sabrina Liberatori, Sibah Hasan, Kristopher Clark, Philip Cohen, Sridhar Vasudevan, Stuart Peirson, Shabaz Mohammed, Vladyslav Vyazovskiy, Russell G Foster, Aarti Jagannath

**Affiliations:** Sleep and Circadian Neuroscience Institute (SCNi), Nuffield Department of Clinical Neurosciences, New Biochemistry Building, University of Oxford, South Parks Road, Oxford, OX1 3QU, U.K.; Departments of Chemistry and Biochemistry, University of Oxford New Biochemistry building, South Parks Road, Oxford, OX1 3QU, U.K.; MRC Protein Phosphorylation and Ubiquitylation Unit, University of Dundee, Sir James Black Centre, Dow Street, Dundee DD1 5EH, U.K.; Department of Pharmacology, University of Oxford, Mansfield Road, Oxford OX1 3QT, U.K.; Department of Physiology, Anatomy and Genetics, University of Oxford, South Parks Road, Oxford OX1 3PT, U.K.

**Author notes:** Correspondence to AJ and RGF.

## Abstract

The sleep and circadian systems act in concert to regulate sleep-wake timing, yet the molecular mechanisms that underpin their interaction to induce sleep remain unknown. Synaptic protein phosphorylation, driven by the kinase SIK3, correlates with sleep pressure, however it is unclear whether these phosphoproteome changes are causally responsible for inducing sleep. Here we show that the light-dependent activity of SIK1 controls the phosphorylation of a subset of the brain phosphoproteome to induce sleep in a manner that is independent of sleep pressure. By uncoupling phosphorylation and sleep induction from sleep pressure, we establish that synaptic protein phosphorylation provides a causal mechanism for the induction of sleep under different environmental contexts. Furthermore, we propose a framework that details how the salt-inducible kinases regulate the synaptic phosphoproteome to integrate exogenous and endogenous stimuli, thereby providing the molecular basis upon which the sleep and circadian systems interact to control the sleep-wake cycle.

## INTRODUCTION

Sleep is a reversible, complex and near ubiquitous behavioural state that results from the interaction of multiple neuronal, hormonal and biochemical circuits (Scammell et al., 2017). Currently the dynamics of sleep are explained by the two-process model, which details two drives that act in concert to regulate sleep timing; the circadian system, known as Process C, and sleep homeostasis, known as Process S (Borbély, 1982; Borbély et al., 2016). Our molecular understanding of Process C has advanced rapidly over the past few decades, and we now know that circadian rhythms, the near 24-hour oscillations in multiple physiological and behavioural processes, are generated by the molecular circadian clock; a complex series of interconnected transcription-translation feedback loops (Takahashi, 2017). This timekeeping mechanism is found in almost every cell of the body, and these clocks are both synchronised with one another and aligned (entrained) to the external environment (Golombek and Rosenstein, 2010; Legates et al., 2014), in a process orchestrated by a master circadian pacemaker located within the suprachiasmatic nuclei (SCN) (Foster et al., 2020; Hastings et al., 2018; Jagannath et al., 2013; Mieda, 2019). Ultimately this allows the temporal optimisation of physiology and behaviour to the varied demands of day and night (Reppert and Weaver, 2002).

In comparison, the exact molecular nature of Process S and the molecular substrates that underpin and control sleep still remain almost entirely unknown. Furthermore, whilst the two-process model provides an accurate behavioural level description of the sleep-wake cycle, the molecular basis of the interaction between the sleep and circadian system that acts to drive sleep has yet to be elucidated (Ode and Ueda, 2020). However, recent studies have highlighted the possible role of the synaptic phosphoproteome in sleep induction and regulation, as synaptic protein phosphorylation has been found to correlate with high levels of sleep pressure, and thus sleep itself. Notably, the salt-inducible kinase, SIK3 is proposed to mediate these synaptic phosphoprotein changes (Brüning et al., 2019; Wang et al., 2018). Whilst providing the foundation for the intriguing hypothesis that the molecular basis of sleep induction may be encoded at the level of the synaptic phosphoproteome, currently these studies cannot establish causation, and instead only demonstrate a correlation between synaptic protein phosphorylation and sleep (Ode and Ueda, 2020). Furthermore, whether sleep-promoting external inputs such as light, that operate independently of sleep pressure and/or sleep history, also impact the synaptic phosphoproteome is entirely unknown. Therefore, in this study we sought to determine whether synaptic protein phosphorylation provides a causal and universal mechanism for the induction of sleep under different environmental contexts.

To address this aim, we focussed our efforts on the salt-inducible kinases, as SIK3 has been proposed to mediate the synaptic phosphoprotein changes observed at times of high sleep pressure (Wang et al., 2018), and we have shown previously that the related kinase SIK1 is light-induced and determines the rate of circadian re-entrainment following nocturnal light exposure as a consequence of a delayed or advanced light/dark cycle (Jagannath et al., 2013). As related kinases have similar substrates, we hypothesised that SIK1 might also regulate this core set of synaptic phosphoproteins to provide information on environmental light to the sleep system, independently from sleep pressure. If true, this would allow us to establish causality by uncoupling synaptic protein phosphorylation and sleep induction from sleep pressure. By utilising a SIK1 kinase-inactive transgenic mouse line, we demonstrate that the light-dependent activity of SIK1 controls the phosphorylation of a subset of the brain phosphoproteome to induce sleep in a manner that is independent of sleep pressure. This allows us to conclude, for the first time, that synaptic protein phosphorylation is causally responsible for sleep induction and provides the molecular substrate by which Process S and Process C interact to regulate the sleep-wake cycle. Furthermore, our data highlights the SIK family as a key regulator of this process that has evolved to buffer the sleep and circadian systems against dynamic environmental and physiological challenges.

## RESULTS

### Light induced SIK1 regulates the induction of sleep in a manner independent from sleep pressure

To examine the hypothesis that the activity of SIK1 induces sleep in response to light independently from sleep pressure, we examined the behavioural phenotype of SIK1 kinase-inactive knock-in (KI) mice, in which the wild type (WT) *Sik1* gene has been replaced with a catalytically inactive version (Darling et al., 2016). We predicted that if SIK1 is induced only after a light/dark cycle shift, as we have previously shown (Jagannath et al., 2013), then SIK1 KI animals would have normal circadian activity and sleep architecture under baseline conditions, but would display abnormal circadian behaviour following nocturnal light exposure that occurs as a consequence of the shifted light/dark cycle (Figure 1A). This is indeed the case. We undertook extensive circadian and sleep phenotyping of WT and SIK1 KI animals and found no major circadian or EEG sleep differences between WT and SIK1 KI mice housed under stable light/dark or constant conditions (Figures S1 and S2). Nocturnal light/CREB signalling, however, resulted in the specific induction of SIK1, and not its related family members SIK2 or SIK3 (Figures S3A-S3I), highlighting that SIK1 exclusively encodes environmental light input.

**Figure 1.**
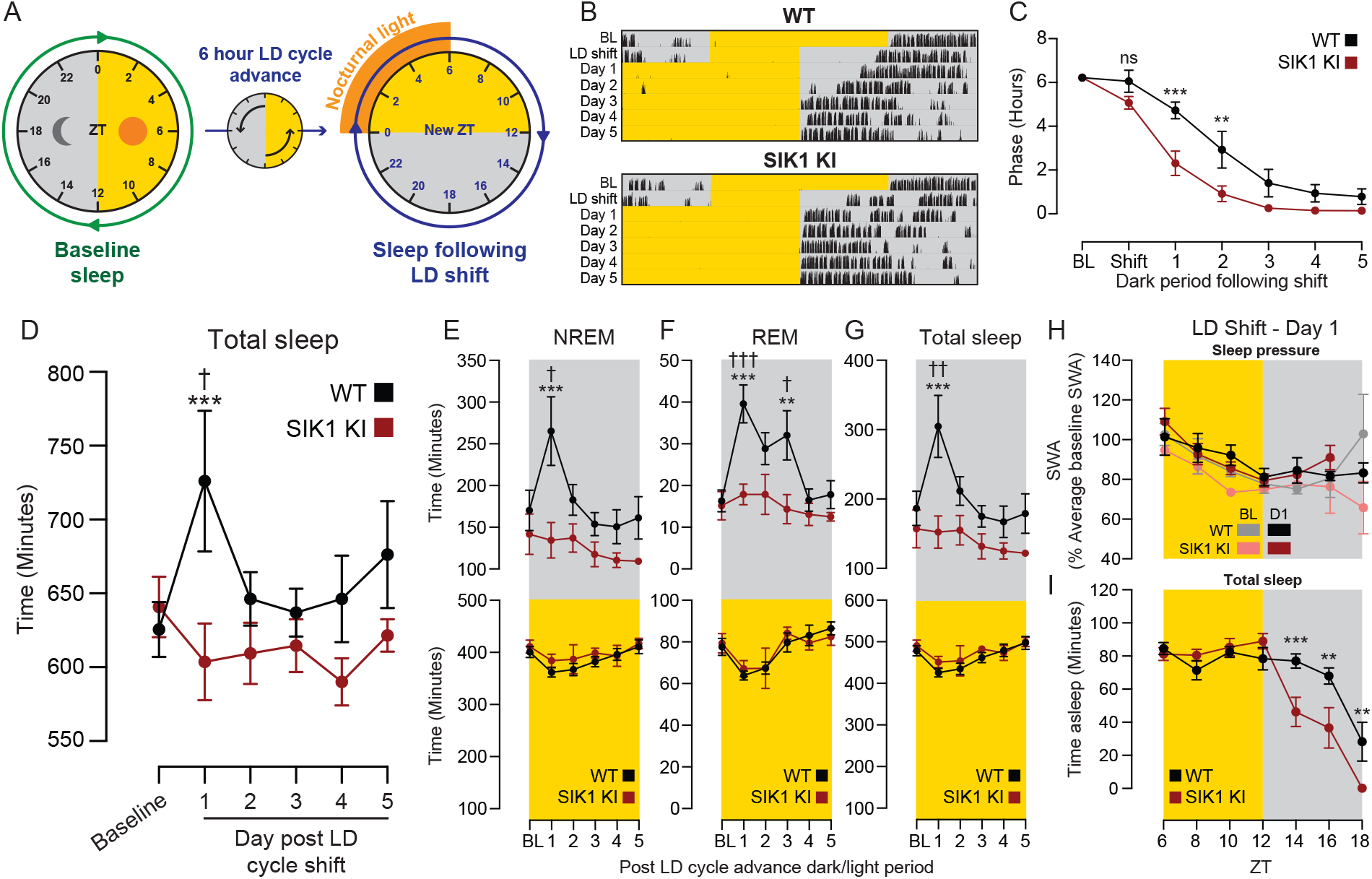
The light-inducible kinase SIK1 regulates the induction of sleep after nocturnal light exposure in a manner independent from sleep pressure. WT and SIK1 KI animals underwent surgery to implant EEG and EMG electrodes and then were housed under a 12:12 LD cycle with continuous sleep recording. (A) Baseline sleep (BL) was recorded, the light/dark (LD) cycle advanced by 6 hours, and sleep and running wheel activity in the days following the LD cycle shift measured. (B) Representative actograms of WT and SIK1 KI animals with baseline, LD cycle shift and days following this shift indicated. (C) Quantification of phase confirmed the SIK1 KI rapid entrainment phenotype in the sleep study cohort. (D) Quantification of total daily sleep time found that WT animals slept significantly more in the first full day after the LD cycle shift, in comparison to the baseline day, whereas SIK1 KI animals did not display an increase in sleep on any day following the LD cycle shift. In comparison to WT animals, SIK1 KI mice did not have increased (E) NREM, (F) REM or (G) total sleep time in the dark period (grey box) of first full day, or any of the subsequent days, following the LD cycle shift. There was no difference in any of the above sleep measures between the genotypes during the light period (yellow box) following the LD cycle shift. There was no difference in (H) the levels of SWA or (I) time spent asleep (both calculated in 2-hour bins) in the 6 hours prior to dark onset of shift day 1 (see B), however (I) SIK1 KI animals spent significantly less time asleep than WT mice 2, 4 and 6 hours after dark onset of shift day 1. Data are mean ± SEM, n = 5-6. Statistical analysis was conducted by two-way ANOVA with Sidak’s correction. ns P > 0.05, ** P < 0.01, *** P < 0.001.

When then subjected to a 6-hour light advance, SIK1 KI animals displayed significantly enhanced circadian behavioural re-entrainment (Figures S3J and S3K). Importantly, this rapid re-entrainment phenotype was entirely light-driven, as during a 6-hour dark advance, the enhanced behavioural shifting of SIK1 KI animals was delayed until after exposure to the first advanced light period (Figures S3L and S3M). SIK1 KI animals also displayed enhanced re-entrainment to a series of light intensities (Figures S3N-S3R) and had greater phase-shifting responses to phase delaying and advancing light pulses (Figures S3S and S3T). Additionally, SIK1 KI animals suppressed wheel-running activity (masked) normally in response to nocturnal light (Figure S3U). Notably, when released into constant darkness 3 days after a 6-hour light advance, their enhanced behavioural re-entrainment was maintained (Figures S3V-S3X), with a significant period shortening in the first 3 days of darkness (Figures S3V, S3W and S3Y). This demonstrates that the re-alignment was due to a shifting of the circadian pacemaker rather than a masking effect or an artefact of the altered light/dark cycle. Collectively, these data demonstrate that SIK1, in an entirely light-dependent manner, directly regulates behavioural re-alignment following a light/dark cycle shift.

We next undertook EEG measurement of sleep in WT and SIK1 KI animals before and after a 6-hour light/dark cycle advance (Figure 1A). The SIK1 KI animals displayed rapid behavioural re-entrainment (Figures 1B and 1C) thereby validating that this cohort would allow us to assess the role of SIK1 in regulating sleep dynamics and architecture nocturnal light exposure. Interestingly, the light/dark cycle shift resulted in an acute and transient induction of sleep in WT mice that was entirely absent in the SIK1 KI animals (Figure 1D). We found that nocturnal light induced both non-rapid eye movement (NREM) and rapid eye movement (REM) sleep in WT mice during the dark period after the light/dark shift, whereas SIK1 KI animals displayed a total lack of sleep induction; NREM, REM and total sleep time remained the same as under baseline conditions (Figures 1E-1G).

Crucially, this lack of sleep induction in SIK1 KI animals was not due to underlying differences in sleep pressure. On the first day following the light/dark cycle shift, slow wave activity (SWA - an electrophysiological metric of sleep need (Wang et al., 2018)) was indistinguishable between WT and SIK1 KI mice (Figure 1H), yet SIK1 KI animals spent significantly less time asleep (Figure 1I). Taken together, our data demonstrate that nocturnal light acts through SIK1 to induce sleep in a manner that is entirely independent of sleep pressure.

### Nocturnal light exposure remodels the brain transcriptome and phosphoproteome

The same data show that the sleep induction we observed in WT animals following a light/dark cycle shift was also not caused by changes in sleep pressure (Figures 1H and 1I). A comparison between the time course of sleep on the baseline day and shift day one found that the WT animals spent significantly more time asleep after dark onset (ZT12) on the first full day following the light/dark shift (Figure 2A). Significantly however, SWA was not different, at any time point, between the baseline day and after the light/dark shift (Figure 2B). Furthermore, whilst the increased sleep seen on shift day one (Figure 2A) resulted in higher levels of cumulative sleep pressure in early part of the dark phase (Figure 2C), the total sleep pressure accumulated at the end of shift day one was the same as baseline, demonstrating that this extra sleep was not driven by the sleep homeostat (Figure 2C). Therefore, the relative increase in sleep after light exposure in WT animals was not accompanied by a change in sleep pressure before, during, or after the sleep induction (Figure 1D).

**Figure 2.**
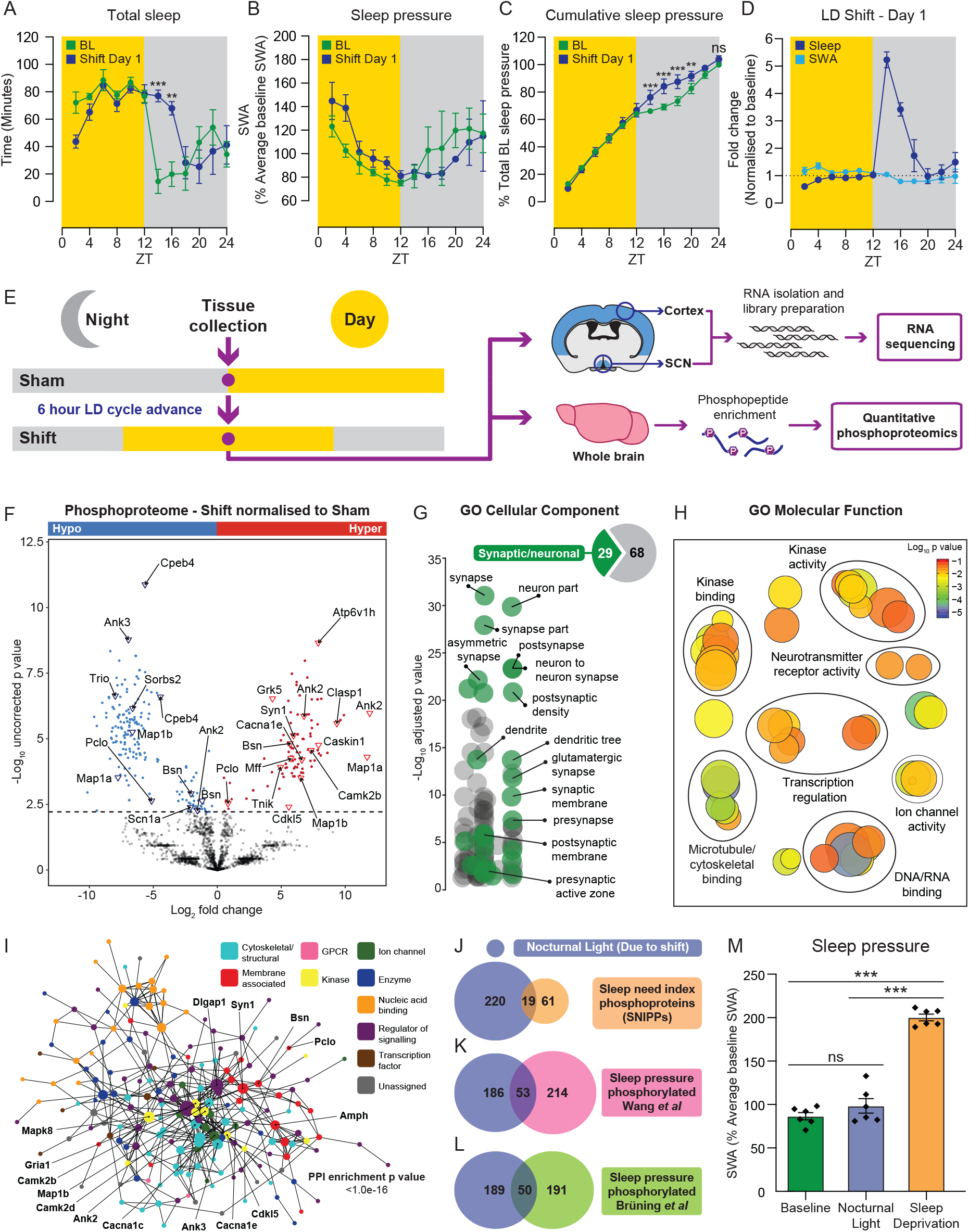
Nocturnal light driven remodelling of the brain phosphoproteome mirrors sleep deprivation, yet occurs independently from sleep need. (A) Total sleep time and (B) average slow wave activity (SWA) in 2-hour bins for the baseline day (green) and first full day after the light/dark cycle advance (blue – Shift Day 1) (C) Cumulative sleep pressure in 2-hour bins, as a percentage total baseline sleep pressure, on the baseline day and shift day 1. (D) Fold change of sleep (dark blue) and SWA (light blue) between shift day 1 and the baseline day. (E) SCN and cortex (RNA sequencing), and whole brain (Quantitative phosphoproteomics) samples were collected either 6 hours into a 6-hour light advance (Shift) or at the same relative time, but without a light shift (Sham). (F) Mass spectrometry analysis of shift and sham WT samples following phosphopeptide enrichment. (G) GO cellular component, (H) GO molecular function pathway analysis, and (I) STRING protein-protein interaction (PPI) analysis of significant phosphopeptides. The overlap between our nocturnal light phosphoproteome and (J) the SNIPPs, or all sleep pressure hyperphosphorylated proteins detailed in (K) Wang *et al*. or (L) Brüning *et al*. (M) SWA 6 hours into the light period on the baseline day, on the first day after the light/dark cycle shift, or after sleep deprivation. Data are (A-D and M) mean ± SEM or (E) mean only. Statistical analysis was conducted by (A-D) two-way ANOVA with Sidak’s correction, by multiple student’s two-tailed t-tests with a two-stage step up FDR correction, or by (M) one-way ANOVA with Tukey’s correction. n = 3-6. ns P > 0.05, ** P < 0.01, *** P < 0.001.

Next, we wanted to determine the molecular mechanism that underpins nocturnal light-induced sleep. As SIK1 regulates CREB-dependent transcription in response to light and protein phosphorylation, we characterised the SCN and cortical transcriptome, and the whole brain phosphoproteome, of WT mice subjected a six-hour light/dark cycle advance (Figure 2E). RNA sequencing of the SCN and cortex six hours into the shifted light/dark cycle identified 537 and 102 genes respectively, that were differentially regulated following nocturnal light exposure (Figure S4 and Table S1). These genes mapped on to molecular functions that were largely distinct between the two tissues, however protein phosphorylation terms, including kinase activity and binding, as well as phosphatase activity, were enriched in both the SCN and cortex following nocturnal light exposure (Figures S4B, S4C, S4E and S4F).

Quantitative phosphoproteomics of whole brains harvested under sham or advanced light/dark cycle conditions (Figure 2E) identified 10,778 phosphopeptides in total (derived from 2730 proteins), of which 324 (mapping to 239 unique proteins) were found to be light regulated (Figure 2F and Table S2). Of these, 217 were hypophosphorylated and 107 were hyperphosphorylated (Figure 2F). GO cellular component analysis found enrichment for pre and postsynaptic proteins (Figure 2G), including microtubule-associated proteins (MAP1A/B), ion channels (SCN1A, CACNA1E), NMDA receptor subunits (GRIN2B), kinases (CAMK2B and CDKL5), synaptic vesicle proteins (SYN1) and presynaptic active zone proteins (BSN, PCLO) (Figure 2F), indicating that these phosphoprotein changes impact upon important neuronal and synaptic functions and processes. Indeed, GO molecular function pathway analysis found overrepresentation of kinase binding and activity, transcriptional regulation, ion channel and neurotransmitter receptor function, and microtubule binding (Figure 2H). STRING protein-protein interaction mapping found that these phosphoproteins form a densely connected interaction network (Figure 2I), and demonstrated that these proteins act together as a highly integrated set to orchestrate synaptic, neuronal and whole brain physiology following nocturnal light exposure.

### The light induced and the sleep need induced phosphoproteomes overlap and independently result in behavioural sleep induction

A large number of these phosphoproteins have been shown previously to be differentially phosphorylated at times of high sleep need. Wang *et al*. document a list of 80 proteins, termed sleep need index phosphoproteins (SNIPPs), whose hyperphosphorylation correlates with increased sleep need(Wang et al., 2018). Notably 19 of these proteins overlapped significantly with the phosphoproteins identified in our study (Figure 2J - Fisher’s exact test p value = 3.5 × 10^−5^). Interestingly, these SNIPPs were curated from a larger list of proteins that were phosphorylated in response to sleep deprivation and/or in the *Sleepy* transgenic mouse line, which has increased daily sleep due to constitutively high SWA (Funato et al., 2016; Honda et al., 2018; Wang et al., 2018). When comparing the intersection of our results with this larger set, we found a highly significant overlap of 53 phosphoproteins (Figure 2K – Fisher’s exact test p value = 1.6 × 10^−9^). In addition, a recent study by Brüning *et al* examining brain protein phosphorylation over the circadian day, and in response to sleep deprivation, also found synaptic protein hyperphosphorylation at times of increased sleep pressure (Brüning et al., 2019). Of these, 50 overlapped with our nocturnal light phosphoproteins (Figure 2L – Fisher’s exact test p value = 1.0 × 10^−9^). Therefore, the light induced and sleep need-induced synaptic phosphoproteomes clearly overlap (Table S3).

However, at the corresponding time where we found altered protein phosphorylation in WT animals, we observed no change in SWA; indeed the level was indistinguishable to SWA under the equivalent baseline condition (Figure 2M), whereas a six-hour sleep deprivation conducted in the same animals as a positive control significantly increased SWA (Figure 2M). This clearly indicated that the changes we observed in protein phosphorylation (Figure 2F) were not caused by increased sleep pressure, but rather by nocturnal light exposure. Collectively, these results suggested to us that synaptic protein phosphorylation in response to light may be causally responsible for sleep induction.

### SIK1 induced changes in the brain phosphoproteome are necessary and sufficient to induce sleep following nocturnal light exposure

As nocturnal light exposure remodels the synaptic phosphoproteome and induces sleep in WT mice, but this sleep induction is missing in SIK1 KI animals, we hypothesised that SIK1 co-ordinates the synaptic phosphoproteome to induce sleep in response to nocturnal light. Therefore, we examined the whole brain phosphoproteome of SIK1 KI animals following the same light/dark cycle shift as used for the sleep analysis (Figure 2E), and anticipated that SIK1 KI animals would have an abnormal phosphoproteome following nocturnal light exposure. Indeed, we found that 99 phosphoproteins (from 88 unique proteins) were differentially regulated between WT and SIK1 KI animals following nocturnal light exposure, with 57 hyper- and 42 hypophosphorylated in SIK1 KI animals in comparison to WT mice (Figure 3A). Many of these phosphoproteins are both nocturnal light and sleep pressure associated (Figures 2F and 2J-2L), including PCLO, BSN, ANK2, AKAP12, SORBS2 and GRK5, and the majority are hypophosphorylated in SIK1 KI animals (Figure 3A and Table S2). Critically, these phosphoprotein differences were not due to underlying differences in SWA architecture in SIK1 KI animals, as they displayed identical SWA levels to WT mice under baseline conditions and during the first day of the shifted light/dark cycle (Figure 1H). GO molecular function pathway analysis of these differential phosphoproteins found overrepresentation of terms similar to those in the WT nocturnal light phosphoproteome (Figures 3B and 2H), demonstrating that these pathways are dysregulated in SIK1 KI animals. Indeed, 35 proteins that were differentially phosphorylated in response to nocturnal light in WT mice displayed a different phosphorylation state in SIK1 KI animals (Figure 3C and Table S3), with many mapping onto recognised elements of light entrainment pathways within the SCN. For example, ADCY5 controls the production of cAMP (Dessauer et al., 2017), ATF2 mediates CREB-dependent gene transcription (Watson et al., 2017), and MAPK8 and MAPK10 regulate light-induced phase shifting (Goldsmith and Bell-Pedersen, 2013; Yoshitane et al., 2012).

**Figure 3.**
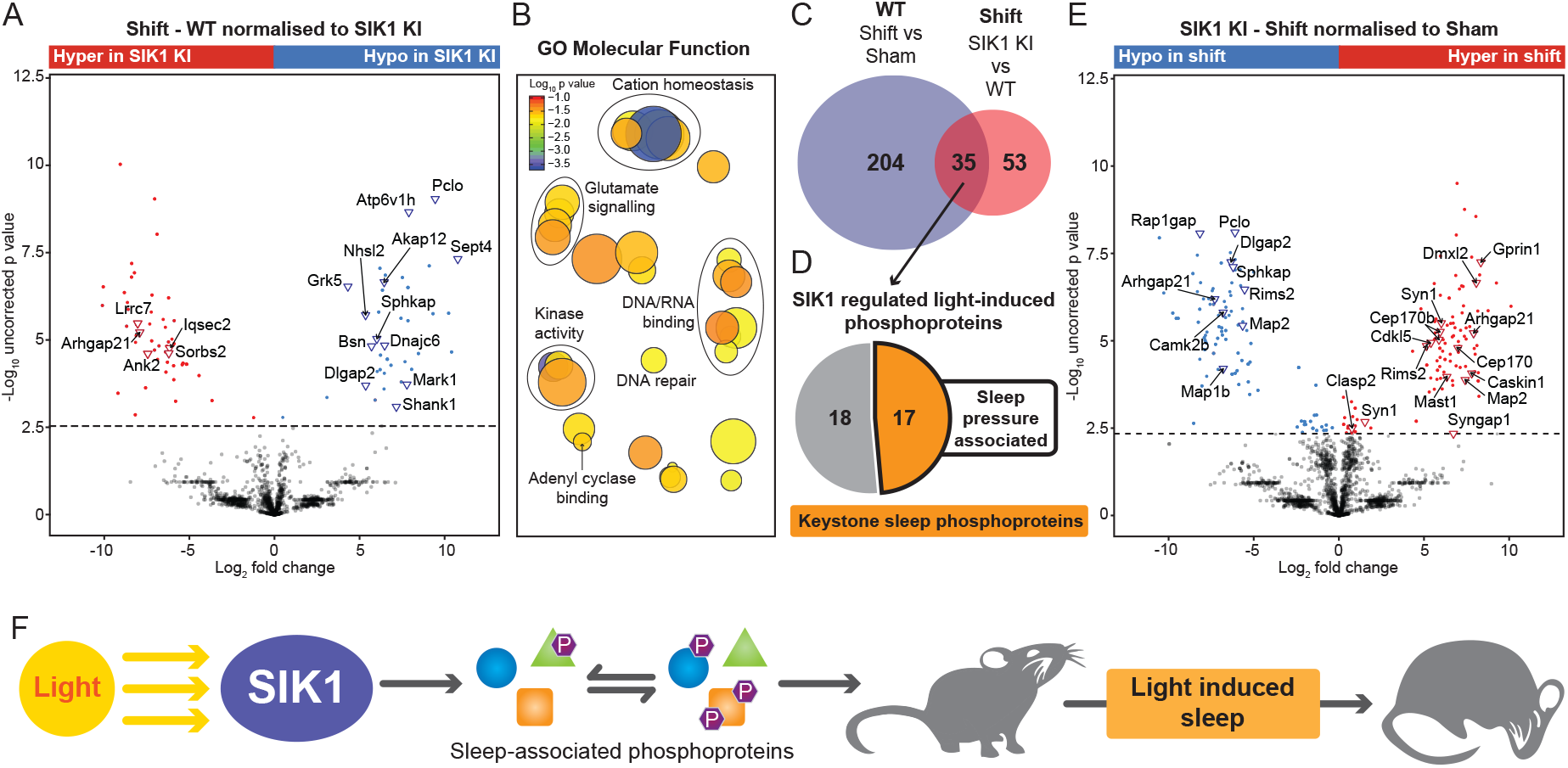
SIK1 induced changes in the brain phosphoproteome are necessary and sufficient to induce sleep following nocturnal light exposure. (A) Phosphopeptide enrichment analysis of whole brain WT and SIK1 KI shift samples. (B) GO molecular function pathway analysis of differential phosphopeptides. (C) Overlap between the WT shift and SIK1 KI differential phosphoproteins, with (D) 17 being previously sleep pressure associated (Termed here keystone sleep phosphoproteins). (E) Phosphopeptide enrichment analysis of SIK1 KI shift vs sham samples. (F) SIK1 controls a subset of the brain phosphoproteome to induce sleep in response to nocturnal light. (A, E) Mean only, multiple student’s two-tailed t-tests with a two-stage step up FDR correction (Q = 0.2).

Notably however, 17 of these SIK1 regulated phosphoproteins have been linked previously to sleep/wake history (Figure 3D and Table S3), and we suggest that these phosphoproteins form the core mechanism that conveys molecular level changes at the synapse to the causal induction of sleep. As such we have termed these keystone sleep phosphoproteins (KSPs). When these are phosphorylated, as is the case in WT mice following nocturnal light exposure, sleep is induced (Figures 1D-1G and 2F). However, when they are not phosphorylated, as is the case in SIK1 KI mice after nocturnal light exposure, sleep is not induced (Figures 1D-1G and 3A). Therefore, SIK1 induced changes in the synaptic phosphoproteome are necessary and sufficient to induce sleep following nocturnal light exposure. Intriguingly, some phosphorylation changes persist, and are even gained, in the absence of a functional SIK1 (Figures 3C, 3E and Table S2), and demonstrates that the phosphoprotein landscape is complex and likely relies on a cascade of kinase and phosphatase activity. Indeed, CAMK2B, CAMK2D, MAPK8, MAPK10, PPP6R1 and PTPN4 displayed differential phosphorylation in our data set, and CAMK2A/B and ERK have been previously shown to regulate sleep duration and dynamics (Mikhail et al., 2017; Tatsuki et al., 2016). Together these results demonstrate that the light-dependent activity of SIK1 controls the phosphorylation of a subset of the brain phosphoproteome to induce sleep in response to nocturnal light, in a manner that is independent of sleep pressure (Figure 3F).

### SIK1 KI animals display normal sleep induction and synaptic protein phosphorylation following acute sleep deprivation

It is clear that SIK1 KI animals do not correctly regulate the phosphorylation of KSPs in response to light and sleep is not induced. In order to confirm causality, we sought to determine whether we could achieve the correct phosphorylation of the KSPs via a different modality in SIK1 KI animals, and whether this would result in normal sleep induction. If so, this would demonstrate that synaptic protein phosphorylation is indeed causal for the induction of sleep. To this end, we utilised the fact that SIK3 activity is preserved in SIK1 KI animals (Darling et al., 2016), and subjected them to six hours of sleep deprivation. In this scenario, SIK3 would drive synaptic protein phosphorylation due to the increased sleep pressure, and therefore if these phosphorylation changes are essential for sleep induction, SIK1 KI animals should display normal sleep architecture and homeostasis and synaptic protein phosphorylation following acute sleep deprivation.

Indeed, quantitative whole brain phosphoproteomics demonstrated that SIK1 KI animals differentially regulated 2289 phosphopeptides (1166 hyperphosphorylated and 1123 hypophosphorylated) following sleep deprivation, with approximately 50% being previously sleep need associated (Figure 4A and Table S4). Many of these were also keystone sleep phosphopeptides (Figures 4A, 3D and Tables S3 and S4). Importantly, this phosphorylation landscape closely mirrored that of the WT animals, with the average phosphopeptide intensities being highly correlated (Pearson r = 0.8125) between WT and SIK1 KI sleep deprived samples (Figure 4B). Indeed, there was a highly significant overlap between the phosphoproteins that displayed differential phosphorylation in WT and SIK1 KI animals, demonstrating that the vast majority of the phosphoprotein landscape is similarly regulated in both genotypes following sleep deprivation (Figure 4C - Fisher’s exact test p value = 1.6 × 10^−48^ and Table S4). In contrast, there was much less similarity between the nocturnal light-driven WT and SIK1 KI phosphoproteomes as their degree of overlap was far smaller (Figure 3C). Critically, 131 (54%) of the Brüning *et al* sleep associated phosphoproteins (Figure 4D), 121 (45%) of the Wang *et al* sleep associated phosphoproteins (Figure 4E), 42 (53%) of the SNIPPs (Figure 4F) and, most importantly, 12 (71%) of the keystone sleep phosphoproteins (Figure 4G) were similarly regulated in both WT and SIK1 KI animals after acute sleep deprivation (Table S4). Therefore, SIK1 KI animals correctly phosphorylate sleep associated phosphoproteins following sleep deprivation.

**Figure 4.**
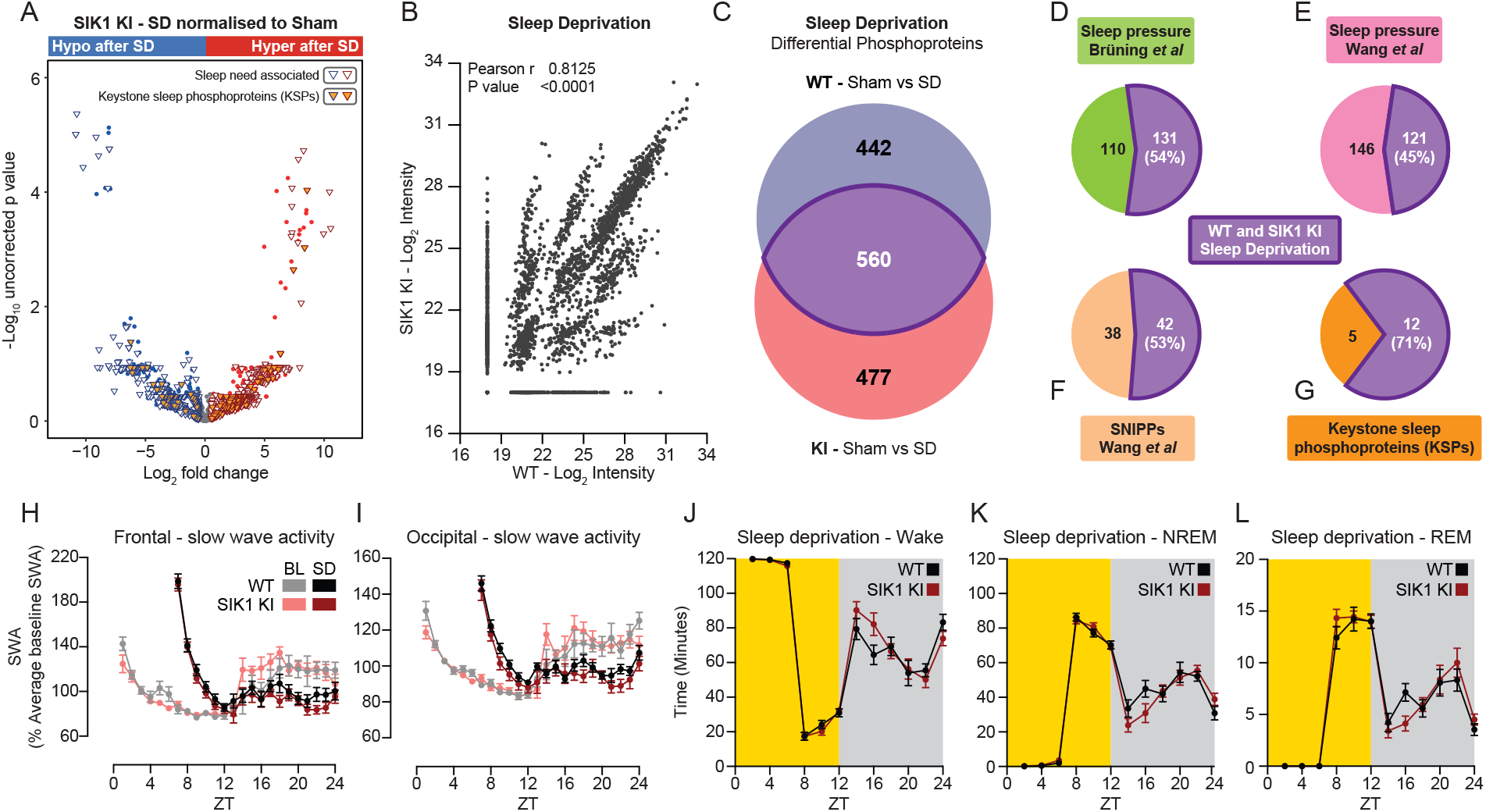
SIK1 KI animals correctly phosphorylate synaptic phosphoproteins following 6 hours of sleep deprivation, which results in normal sleep and slow wave activity architecture. WT and SIK1 KI animals were housed under a 12:12 LD cycle with continuous sleep recording. Sleep deprivation was then conducted from ZT0 – ZT6. Mass spectrometry following phosphopeptide enrichment was conducted on WT and SIK1KI whole brain samples following SD. (A) Volcano plot of differential phosphopeptides (blue – hypophosphorylated after SD; red – hyperphosphorylated after SD) in SIK1 KI mice following SD. All previously sleep need associated phosphoproteins (open triangles) and keystone sleep phosphoproteins (orange filled triangles – as detailed in Figure 3D) are highlighted. (B) Correlation analysis of phosphopeptide abundance between WT and SIK1 KI SD samples. (C) Overlap of WT and SIK1 KI differential phosphopeptides with the same directionality. The number of WT and SIK1 KI SD differential phosphopeptides that are present in (D) the sleep pressure associated phosphoproteins from Brüning *et al*. or (E) *Wang et al*., (F) SNIPPs from Wang *et al*. and (G) the keystone sleep phosphopeptides. There was no difference in the levels of (H) frontal and (I) occipital SWA under baseline conditions (BL), or following SD, between the genotypes, and the amount of (J) wake, (K) NREM and (L) REM, in 2-hour bins, did not differ between WT and SIK1 KI animals following SD. Data are (A,B) mean only or (H-L) mean ± SEM or. n = (H-L) 12-14 and (A-G) 3. For (H-L) statistical analysis was conducted by two-way ANOVA with Sidak’s multiple comparisons correction. ns P > 0.05.

This was accompanied by subsequent normal sleep induction and architecture as in comparison to WT mice, SIK1 KI animals exhibited no discernible difference in sleep homeostasis following sleep deprivation; they accumulated and dissipated SWA identically (Figures 4H and 4I) and displayed completely normal rebound sleep architecture (Figures 4J-4L and Figure S5). Taken together, these results demonstrate that synaptic protein phosphorylation is necessary and sufficient, and therefore causal, for the induction of sleep. Furthermore, they advance our basic understanding of the regulatory mechanisms underpinning sleep and highlight the salt-inducible kinase family as a key regulator of synaptic protein phosphorylation and sleep induction.

## DISCUSSION

In this study we utilised a SIK1 kinase-inactive mouse line to uncouple phosphorylation and sleep induction from sleep pressure and determine that synaptic protein phosphorylation provides a causal mechanism for the induction of sleep under different environmental contexts. In fact, the usage of a loss-of-function model was central to our discovery that SIK1 controls synaptic protein phosphorylation and sleep induction only after exposure to nocturnal light. Whilst previous studies have proved seminal in demonstrating the role of salt-inducible kinases in regulating sleep, they have all used gain-of-function mutants (Funato et al., 2016; Honda et al., 2018; Park et al., 2020; Wang et al., 2018), which cannot explain how the kinase is endogenously regulated, and therefore have likely missed exactly when and how the SIKs act to regulate sleep. Indeed, a previous study examining the role of SIK1 in sleep found that the *Sik1*^*S577A*^ gain-of-function mutant displayed increased sleep duration under stable light/dark conditions (Park et al., 2020). This is not surprising as SIK1 is able to phosphorylate the same targets as SIK2 or SIK3 when activated, however SIK1 is exclusively induced by light, which could not be captured in the study above (Park et al., 2020). Instead, we show here that SIK1 inactive mutant animals had normal baseline sleep, and only displayed sleep abnormalities following nocturnal light exposure.

We also demonstrate here that the light-dependent activity of SIK1 is necessary and sufficient to coordinate circadian and behavioural realignment after a light/dark cycle shift. A group of 35 proteins were identified, whose phosphorylation is controlled by SIK1 in response to nocturnal light, and we propose that these phosphoproteins play a major role is adjusting physiology and behaviour to the astronomical day. Indeed, many of these proteins are regulatory elements of light-dependent entrainment within the SCN. For example, ATF2 is a transcription factor that heterodimerises with CREB to mediate gene transcription by binding promoter localised cAMP response elements (Watson et al., 2017), and the membrane bound adenylyl cyclase ADCY5 controls the production of cAMP (Dessauer et al., 2017). Alongside, AKAP12 regulates the cellular localisation of PKA (Sanderson and Dell’Acqua, 2011) and this cAMP/PKA/CREB pathway is central for mediating light-induced gene expression and shifting of the molecular circadian clock (O’Neill et al., 2008). Furthermore, we found that SIK1 also controlled the phosphorylation of MAPK8 and MAPK10; two kinases known to regulate phase shifting in response to nocturnal light (Goldsmith and Bell-Pedersen, 2013; Yoshitane et al., 2012).

Importantly however, we detail 17 phosphoproteins identified as both SIK1 and sleep need regulated, termed keystone sleep phosphopeptides, that we believe lie at the centre of the mechanisms that convey molecular level changes at the synapse to the causal induction of sleep. Indeed, these proteins are known to regulate synaptic organisation, presynaptic neurotransmitter release and postsynaptic signalling. Presynaptically, AMPH plays a crucial role in controlling synaptic vesicle recycling (Di Paolo et al., 2002), and the presynaptic active zone proteins BSN and PCLO are essential for regulating synaptic vesicle docking, fusion and neurotransmitter release (Gundelfinger et al., 2016). Postsynaptically, many of the sleep-inducing phosphoproteins are core components of the postsynaptic density (PSD) and regulate downstream neurotransmitter signalling. ANK2 acts as a scaffolding protein that regulates synaptic stability (Bulat et al., 2014; Koch et al., 2008), whereas DLAGP2 and SORBS2 are both PSD scaffolding proteins that regulate excitatory neurotransmission by controlling the turnover and trafficking of AMPA glutamate receptors (Rasmussen et al., 2017; Zhang et al., 2016). Alongside, IQSEC2 (a guanine nucleotide exchange factor) also plays a critical role in glutamate receptor trafficking and regulates synaptic cytoskeletal organisation to impact downstream neurotransmitter signalling (Brown et al., 2016; Um, 2017). Interestingly IQSEC2 has been shown to interact with PSD-95, a scaffolding protein that complexes with HOMER1A to tether glutamate receptors to the PSD (Clifton et al., 2019; Dosemeci et al., 2007; Sakagami et al., 2008). As *Homer1a* is upregulated in response to sleep deprivation (Maret et al., 2007), and serves to drive excitatory synaptic downscaling during sleep (Diering et al., 2017; Martin et al., 2019), this suggests that HOMER1A activity may also contribute to light induced sleep. Additionally, LRRC7, a major constituent of the PSD, controls Ca^2+^ channel flux at excitatory synapses (ANK2 has also recently been found to have a similar function (Choi et al., 2019)) and interestingly, LRRC7 deficient mice have been found to have abnormal sleep behaviour (Carlisle et al., 2011). Similarly, both CAMK2B and CACNA1E were phosphorylated in response to nocturnal light. CAMK2B is a principal regulator of neurotransmitter release and synaptic function (and notably its dendritic localisation is controlled by LRRC7 (Jiao et al., 2011)), whilst CACNA1E forms one of the core Ca^2+^ channels involved in synaptic neurotransmission (Kamp et al., 2005), and strikingly the genetic ablation of either protein results in abnormal sleep length duration (Siwek et al., 2014; Tatsuki et al., 2016). Therefore, we conclude that synaptic protein phosphorylation, in response to either nocturnal light or increasing sleep pressure, controls synaptic organisation and function to lead to the induction of sleep.

Furthermore, we suggest that Process S and Process C, the fundamental components of the two-process model of sleep (Borbély, 1982; Borbély et al., 2016), converge upon the synaptic phosphoproteome through the action of the salt-inducible kinases. Not only does this provide a molecular mechanism by which the sleep and circadian systems interact, this highlights the SIK family as central to this process (Funato et al., 2016; Honda et al., 2018; Park et al., 2020; Wang et al., 2018). Here we propose a model of sleep regulation, whereby SIK1 and SIK3 both control the phosphorylation of a core group of synaptic proteins (Figure 5). Crucially however, their activity is induced by entirely different stimuli, with SIK1 activity dependent on light and SIK3 activity dependent on sleep pressure. Importantly, this model also explains how SIK1 KI animals display normal sleep and circadian behaviour under stable entrainment and after sleep deprivation; SIK1 KI mice have a fully functional copy of SIK3 (Darling et al., 2016). Additionally, the fact that the *Sik3 Sleepy* mutant has been reported to have a normal circadian system (Funato et al., 2016) also supports the model we propose, as SIK1 is intact in *Sleepy* mutant mice. Overall, this framework advances our basic understanding of the regulatory mechanisms underpinning sleep by highlighting how the salt-inducible kinases have evolved to perform distinct – but complementary – roles, to both buffer and integrate the sleep and circadian systems in response to dynamic environmental and physiological challenges.

**Figure 5.**
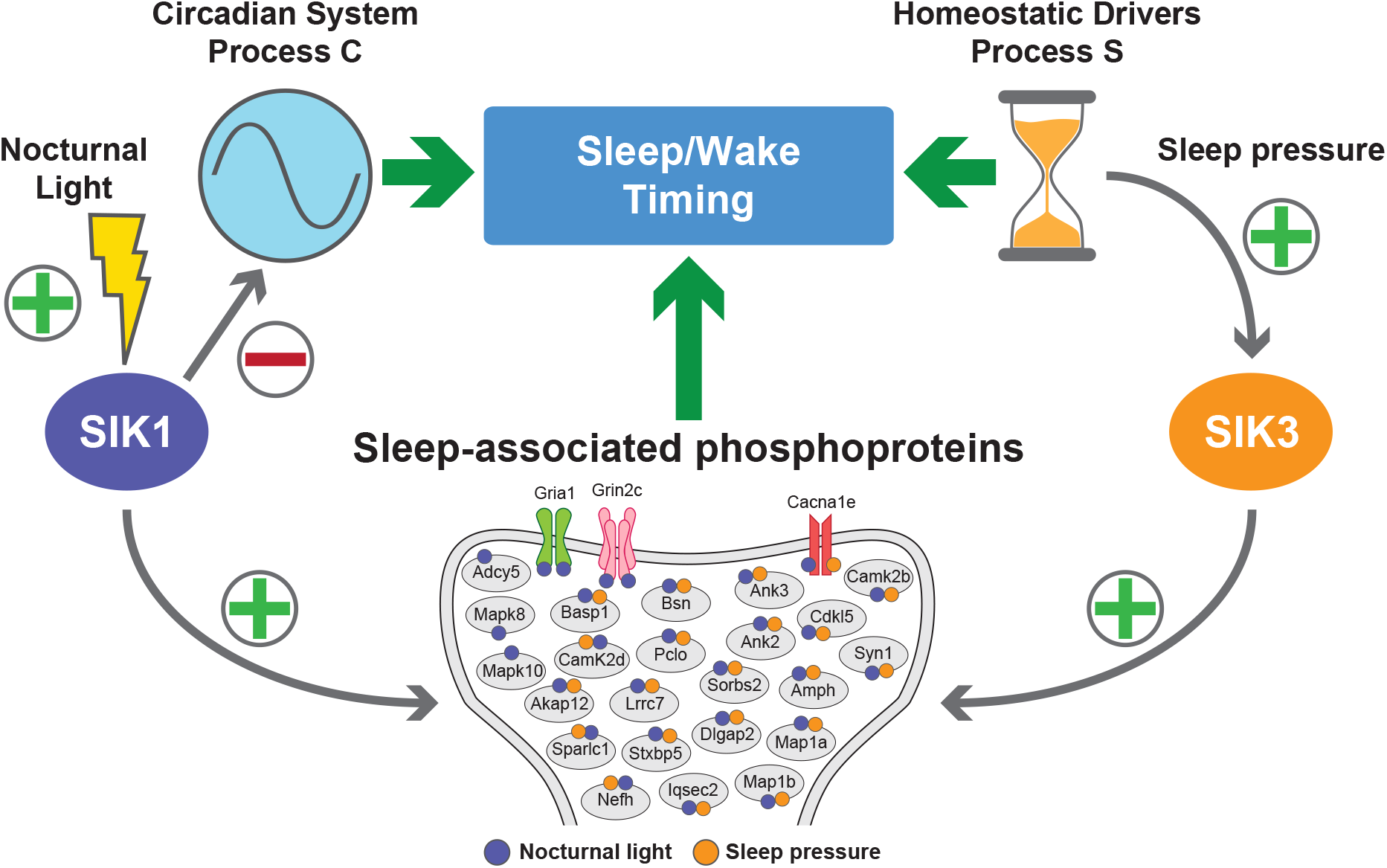
The salt-inducible kinases SIK1 and SIK3 both induce sleep by regulating the phosphorylation status of a core group of synaptic proteins, but under entirely different contexts. We propose a model whereby SIK1 and SIK3, two members of the same kinase family, both induce sleep by regulating the phosphorylation of a core group of synaptic phosphoproteins. Crucially however, their activity is induced by entirely different stimuli, with SIK1 activity dependent on nocturnal light and SIK3 activity dependent on sleep pressure. This model advances our basic understanding of the regulatory mechanisms underpinning sleep by suggesting that Process S and Process C, the fundamental components of the two-process model of sleep, converge upon the synaptic phosphoproteome through the action of the salt-inducible kinases thereby providing a molecular mechanism by which these systems interact. Furthermore, this demonstrates how these kinases have evolved to perform distinct – but complementary – roles, to buffer the sleep and circadian systems in response to environmental and physiological challenge.

## ACKNOWLEDGEMENTS

This work was supported by the following sources of funding: BB/N01992X/1 David Phillips fellowship from the BBSRC to AJ, MRC_MR/K000985/1 to PC, and WT106174/Z/14/ZMA from the Wellcome Trust to RGF.

## AUTHOR CONTRIBUTIONS

LT, TP, BJ, and AJ conducted the experiments, LT, PKR and SW analysed data, KC and PC generated transgenic animals, AJ and RGF supervised the study with SP, SV and VV. In addition, SM oversaw proteomics studies in which SL participated, VV oversaw sleep EEG experiments in which SH participated. LT, RGF and AJ wrote the manuscript with input from all authors.

## COMPETING INTERESTS

None to declare.

## MATERIALS AND METHODS

### RESOURCE AVAILABILITY

#### Lead Contact

Further information and requests for resources and reagents should be directed to and will be fulfilled by Aarti Jagannath

#### Materials Availability

This study did not generate new unique reagents.

#### Data and Code Availability

The datasets generated during this study will be deposited to the appropriate public repositories upon acceptance.

### EXPERIMENTAL MODEL AND SUBJECT DETAILS

#### In vitro cell culture studies

Human U2OS cells and murine NIH3T3 cells were obtained from ATCC (ATCC^®^ HTB-96^™^ and ATCC^®^ CRL-1658^™^ respectively) and cultured in complete DMEM (DMEM supplemented with 10% FCS and 1% penicillin/streptomycin) at 37°C, 5% CO_2_. Cells were lifted by incubation with TrypLE express for 5 mins, diluted with complete DMEM and counted by trypan blue exclusion. Cells were either passaged into new tissue culture flasks, or plated into multi well plates for *in vitro* experiments.

#### In vivo animal studies

C57BL/6-Sik1^tm2853(T182A)Arte^ animals (as previously described in (Darling et al., 2016)) were used throughout this study and are herein referred to as SIK1 knock-in (SIK1 KI) mice. In these animals, the endogenous WT SIK1 gene has been replaced with a mutated version via homologous recombination, which results in the production of a SIK1 protein where threonine 182 (found in the activation loop of SIK1 and its phosphorylation by LKB1 is known to be critical for SIK1 activity) is mutated to alanine, rendering SIK1 catalytically inactive. The WT animals used in this study were littermate and age matched controls. All studies were conducted using mice over 8 weeks of age and unless otherwise indicated, animals were housed in groups with *ad libitum* access to food and water under a 12:12 hour light dark cycle (100 lux from white LED lamps). Animals were randomly assigned to experimental groups. Both male and female animals were used in this study, with the gender used indicated in the corresponding figure legend. All animal procedures were conducted in accordance with the UK Home Office regulations (Guidance on the Operation of Animals (Scientific Procedures Act) 1986) and the University of Oxford’s Policy on the Use of Animals in Scientific research, taking into account the principles of the 3Rs.

### METHOD DETAILS

#### Circadian wheel-running activity monitoring

WT and SIK1 KI animals were individually housed with *ad libitum* access to food and water and circadian running wheel activity measured under a 12 hour light/12 hour dark (100 lux - 12:12 LD) schedule, constant light (100 lux – LL) or constant darkness (DD). White LEDs were used as the light source. Activity data was collected using the ClockLab software package (Actimetrics, Wilmette, IL, USA).

#### Light/dark cycle advance protocols

WT and SIK1 KI animals were individually housed and circadian wheel-running behaviour was measured under a 12:12 LD cycle (100 lux). Following stable behavioural entrainment, either the light or dark phase was advanced by 6 hours and wheel-running activity measured until the mice have re-entrained to the new LD cycle. For the repeated advance protocol, WT and SIK1 KI animals were initially housed under a 12:12 LD cycle with the light intensity set to 100 lux. Following stable entrainment, either the light or dark phase was advanced by 6 hours and wheel-running activity measured. After mice had re-entrained to the new LD cycle, the light intensity was reduced and then the 6-hour shift repeated. This procedure was conducted at 100, 30, 10, 3 and 1 lux. For analysis, phase was calculated as time of activity onset minus time of dark onset.

#### Light pulse and phase shifting protocols

WT and SIK1 KI animals were individually housed and circadian wheel-running behaviour was measured under a 12:12 LD cycle (100 lux). Following stable behavioural entrainment, animals were released into DD and then received 30 minute 100 lux light pulses, timed to fall between CT14-17. The phase angles before and after the light pulse were calculated and the resultant phase shift calculated. As per convention, phase delays are negative and phase advances positive. In order to calculate light induced behavioural masking during the light pulse, the activity counts in the 30 minutes prior to the light pulse and during the 30 minute light pulse were calculated. Only pulse episodes where the 30 minutes prior to the light pulse had non-zero activity counts were included in the analysis.

#### Cell culture and stimulation

Human U2OS cells and murine NIH3T3 cells were obtained from ATCC (ATCC^®^ HTB-96^™^ and ATCC^®^ CRL-1658^™^ respectively) and cultured in complete DMEM (DMEM supplemented with 10% FCS and 1% penicillin/streptomycin) at 37°C, 5% CO_2_. Cells were lifted by incubation with TrypLE express for 5 mins, pelleted, diluted with complete DMEM and counted by trypan blue exclusion, and then plated into 24 well plates (1×10^5^ cells/well in 500 μl) and left overnight at 37°C, 5% CO_2_. The cells were then stimulated with either vehicle (0.5% DMSO), forskolin (10 μM) or dexamethasone (200 nM) for 1 hour at 37°C, 5% CO_2_. Following treatment, the cells were washed three times with 1 ml of PBS, the medium replaced with complete DMEM and then at the time points indicated the cells were lysed by the addition of 300 μl RLT buffer. The lysate was then stored at -80°C until RNA extraction was performed.

#### RNA extraction, cDNA synthesis and RT-PCR

For cells lysed in RLT buffer, RNA was extracted using the RNeasy Plus Mini Kit following the manufacturer’s instructions. For tissue RNA extraction, the tissue was firstly mechanically disrupted in 100 μl of Trizol. The sample was then made up to 500 μl of Trizol, 100 μl chloroform added and then thoroughly mixed. Following a 5 min incubation at RT, the sample was then centrifuged for 15 min at 15,000 xg, 4°C. The clear top layer was then carefully collected, mixed with an equal volume of 70% ethanol and RNA extracted using the RNeasy Plus Mini Kit following the manufacturer’s instructions. RNA concentration and quality was determined using a NanoDrop ND-1000 spectrophotometer and cDNA synthesized using the qScript cDNA synthesis kit. RT-PCR was then conducted using the Quantifast SYBR Green PCR Kit and a StepOnePlus thermal cycler (Applied biosystems) with the following thermal profile: 95°C for 5 mins and then 40 cycles of 95°C for 10s, 60°C for 30s and 72°C for 12 s. Quantification of transcript levels was conducted using the relative standard curve method, for comparing within a gene, or the 2^-ΔCt method for comparison across genes. Primer sequences used can be found in Table S5.

#### RNA sequencing library preparation

Following total RNA extraction, RNA sequencing libraries were prepared using the Illumina TruSeq Stranded Total RNA library prep gold kit following the manufacturer’s instructions. Briefly, 150 ng of total RNA was depleted of ribosomal and mitochondrial ribosomal RNA, cleaned up using RNA clean XP beads (Beckman Coulter, High Wycombe, United Kingdom) and then fragmented. Next, first and second stand cDNA synthesis was conducted, and the resultant cDNA purified using AMPure XP beads (Beckman Coulter) and adenylated at the 3’ end. Illumina indexing adapters were then ligated to the cDNA, the fragments purified and then enriched by PCR. Following amplification, the libraries were purified and then their concentration determined using the KAPA Library Quantification Kit for Illumina Platforms following the manufacturer’s instructions. The libraries were then diluted to 4 nM and pooled in equal volumes prior to sequencing. Paired end RNA sequencing was then conducted using the NextSeq 550 and a Nextseq 500/500 v2 75 cycle kit, with the library loaded at 1.8 pM.

#### RNA sequencing data analysis

The raw reads were initially processed to remove adapter sequences and trim low quality ends. Reads were then mapped to the mouse reference genome (build GRCm38.98) using HISAT2 and gene counts generated using featureCounts (multi-mapping and multi-overlapping reads allowed). Prior to analysis, genes below the minimum expression threshold (total counts less than 10 across all samples) were removed and differential gene expression analysis conducted using DESeq2. Genes that had an uncorrected p value of <0.01 were taken to be statistically significant. For pathway analysis, significant differential gene lists were analysed using Enrichr (Kuleshov et al., 2016) or g:Profiler (Raudvere et al., 2019) and then GO terms reduced and visualised using Revigo (Supek et al., 2011).

#### EEG and EMG electrode implantation

To continuously monitor sleep/wake states, custom electroencephalogram (EEG) and electromyography (EMG) headmounts were implanted as previously described. Briefly, surgical procedures were conducted under aseptic conditions and isoflurane anaesthesia (2-3% for induction, 1-2% for maintenance), with animals head-fixed in a stereotactic frame (David Kopf Instruments, California). Analgesia was administered immediately before the start of surgery (Metacam 1–2 mg/kg s.c. and Vetergesic 0.08 mg/kg s.c.). Screw electrodes, mounted to an 8-pin surface mount connector (8415-SM, Pinnacle Technology Inc, Kansas), were implanted in the frontal (motor area, anteroposterior +2 mm, mediolateral 2 mm) and occipital (visual area, V1, anteroposterior −3.5 to −4 mm, mediolateral 2.5 mm) cortical areas and a reference electrode was implanted above the cerebellum. Two stainless-steel wires were implanted on each side of the nuchal muscle to record EMG and the entire headmount was then fixed to the skull using RelyX Unicem 2 dental cement (3M, Bracknell, UK). Following surgery animals were provided with thermal support, administered saline (0.1 mL/20 g s.c.), and then closely monitored, with analgesia provided if necessary (Metacam 1-2 mg/kg orally), during a 2-week recovery period. For sleep state recording, animals were individually housed in custom clear plexiglass cages (20.3 × 32 × 35 cm) kept in ventilated, sound-attenuated Faraday chambers (Campden Instruments, Loughborough, UK), under a 12:12 LD cycle (100 lux) with *ad libitum* access to food and water. For sleep deprivation, animals were kept awake for 6 hours between ZT0 and ZT6 by regularly providing various novel objects to elicit exploratory behaviour.

#### Sleep signal processing, data acquisition and vigilance state scoring

EEG, EMG and running wheel data was acquired using the Multi-channel Neurophysiology Recording System (Tucker-Davis Technologies, Alachua, FL). The EEG and EMG signals were filtered between 0.1 and 100 Hz, amplified using a PZ5 NeuroDigitizer preamplifier (Tucker-Davis Technologies) and then collected at a sampling rate of 256.9 Hz. Running wheel data (infra-red beam breaks by the running wheel rungs) was collected continuously. The EEG and EMG data were then resampled offline at 256 Hz, converted to .txt files using custom Matlab scripts and then converted to European data format (EDF) using Neurotraces software. Vigilance states were then assigned by manual inspection of consecutive 4 second epochs using the Sleepsign software (Kissei Comtec Co., Nagano, Japan). Vigilance states were classified as wake (low amplitude, high frequency EEG with associated EMG signal), non-rapid eye movement sleep (NREM – high amplitude, low frequency EEG signal with slow waves present) or REM sleep (low amplitude, high frequency theta EEG signal). Brief awakenings were defined as wake periods lasting only 4 epochs or less. Epochs that contained artefactual signals (resulting from contamination by eating, drinking, or gross movement) were scored as such so that they could be excluded from the appropriate analyses. EEG power spectra were then produced using a Fast Fourier Transform routine for all 4-second epochs, with a 0.25-Hz resolution. SWA activity is calculated as the average spectral power between 0.5-4 Hz of all NREM epochs in the desired time bin and expressed as a percentage of the average spectral power between 0.5-4 Hz of all NREM epochs over the entire baseline day.

#### Tissue total protein lysate preparation

WT and SIK1 KI animals were housed under a 12:12 LD cycle (100 lux) and allowed to stably entrain. For sham samples, animals were sacrificed at ZT0 in complete darkness and for the shift samples, the light phase was advanced by 6 hours and the animals sacrificed 6 hours into the advanced light period. Their brains were removed and immediately flash frozen. Whole brain protein extraction was performed by lysing the entire brain in 5 ml tissue lysis buffer (8 M urea, 50 mM HEPES, 0.5% NaDOC, pH 8.5, supplemented with protease and phosphatase inhibitor tablets (Roche)) using a dounce homogeniser (Sigma Aldrich, UK). The samples were then cleared by centrifugation for 20 min at 20,000 xg at 4°C, and protein concentration determined using a BCA protein assay kit (Thermo Fisher scientific, Loughborough, UK) following the manufacturer’s protocol.

#### Mass spectrometry sample preparation and phosphopeptide enrichment

Mass spectrometry sample preparation and phosphopeptide enrichment was conducted as in (Wang et al., 2018), with a few modifications. One milligram of total protein was reduced (10 mM TCEP), alkylated (50 mM CAA) and then digested with Lys-C for 2 h (1:100 enzyme:substrate ratio). The sample was then diluted to 2 M urea using 50 mM HEPES, pH 8.5 and then further digested with trypsin (1:50 enzyme:substrate ratio) overnight at room temperature. After adding TFA to 1% v/v, the samples were centrifuged at 21,000 xg for 10 min at RT, desalted using Oasis desalting columns (Waters) and then dried by vacuum centrifugation. Phosphopeptide enrichment was conducted using titanium dioxide (TiO_2_) beads. Dried peptides were resuspended in binding buffer (65% AcN, 2%TFA) and supplemented with 1 mM KH_2_PO_4_, before being added to TiO_2_ beads that had been washed twice with washing buffer (65% AcN, 0.1%TFA). The samples were incubated for 30 min at RT with constant agitation and then washed twice with washing buffer. The bound phosphopeptides were eluted with 40 μl elution buffer (50% AcN, 14% NH_4_OH pH∼11) and then dried by vacuum centrifugation. The phosphopeptides were then resuspended in 50 mM HEPES pH 8.5, acidified by the addition of TFA to ∼2% v/v, desalted using Oasis desalting columns (Waters) and then dried by vacuum centrifugation.

#### Mass spectrometry data acquisition, processing and analysis

Samples were analysed on an Ultimate 3000 ultra-HPLC system (Thermo Fisher Scientific) and electrosprayed directly into a QExactive mass spectrometer (Thermo Fisher Scientific). They were initially trapped on a C18 PepMap100 pre-column (300 μm inner diameter x 5 mm, 100Å, Thermo Fisher Scientific) in solvent A (0.1% [vol/vol] formic acid in water). The peptides were then separated on an in-house packed analytical column (75 μm inner diameter x 50cm packed with ReproSil-Pur 120 C18-AQ, 1.9 μm, 120 Å, Dr. Maisch GmbH) using a 2h linear 15%-35% [vol/vol] acetonitrile gradient and a flow rate of 200 nl/min. Full-scan mass spectra were acquired in the Orbitrap (scan range 350-1500 m/z, resolution 70000, AGC target 3×10^6^, maximum injection time 50 ms) in a data-dependent mode. After the mass spectrum scans, the top 20 most intense peaks were selected for higher-energy collisional dissociation fragmentation at 30% of normalized collision energy. Higher-energy collisional dissociation fragmentation spectra were also acquired in the Orbitrap (resolution 17500, AGC target 5×10^4^, maximum injection time 120 ms) with first fixed mass at 180 m/z. Peptide identification and quantitation was then performed using MaxQuant (v1.6.3.4). Data was searched against the mouse Uniprot database (January 2017) and a list of common contaminants provided by the software. The search parameters for the Andromeda search engine were: full tryptic specificity, allowing two missed cleavage sites, fixed modification was set to carbamidomethyl (C) and the variable modification to phosphorylation (STY), acetylation (protein N-terminus) and oxidation (M). Match between runs was applied. All other settings were set to default, leading to a 1% FDR for protein identification. For pathway analysis, significant differential protein lists were analysed using Enrichr (Kuleshov et al., 2016) or g:Profiler (Raudvere et al., 2019) and then GO terms reduced and visualised using Revigo (Supek et al., 2011). Protein-protein interaction (PPI) mapping was performed using STRING (Szklarczyk et al., 2019).

### QUANTIFICATION AND STATISTICAL ANALYSIS

Circadian analysis was conducted using the Clocklab software package (Actimetrics Wilmette, IL, USA). Circadian period (Tau) was calculated using a Chi-squared periodogram. The average daily active period (Alpha) is represented in degrees and is corrected for individual circadian period. Bout analysis was conducted using an activity threshold of 4 counts per minute with a maximum permissible gap of 3 minutes. Data visualization and statistical analysis was conducted using GraphPad Prism (Version 8, La Jolla, CA) or RStudio (Version 1.1.447, Boston, MA). All data are expressed as mean + or ± SEM, and n represents the number of independent animals or replicates per group, as detailed in each figure legend. For comparisons between two groups only, a two-tailed Student’s t-test was applied. For multiple comparisons a two-way ANOVA with Sidak’s multiple comparisons correction was used. A p value of <0.05 was taken to be statistically significant. For phosphopeptide analysis following the light/dark cycle advance, log_2_(abundance) values were used for statistical testing. Phosphopeptide differences between groups were assessed using multiple two-tailed unpaired student’s t tests followed by FDR correction using the Benjamini, Krieger and Yekutieli two-stage step up method (Q = 0.2). A Q value of > 0.2 was considered statistically significant. For phosphopeptide analysis following sleep deprivation, differences between groups were assessed by log_2_ fold change. A log_2_ fold change of > 0.5 or < -0.5 was considered differential. The exact test used is specified in the corresponding figure legend.

---

**Figure S1. Related to Figure 1.**
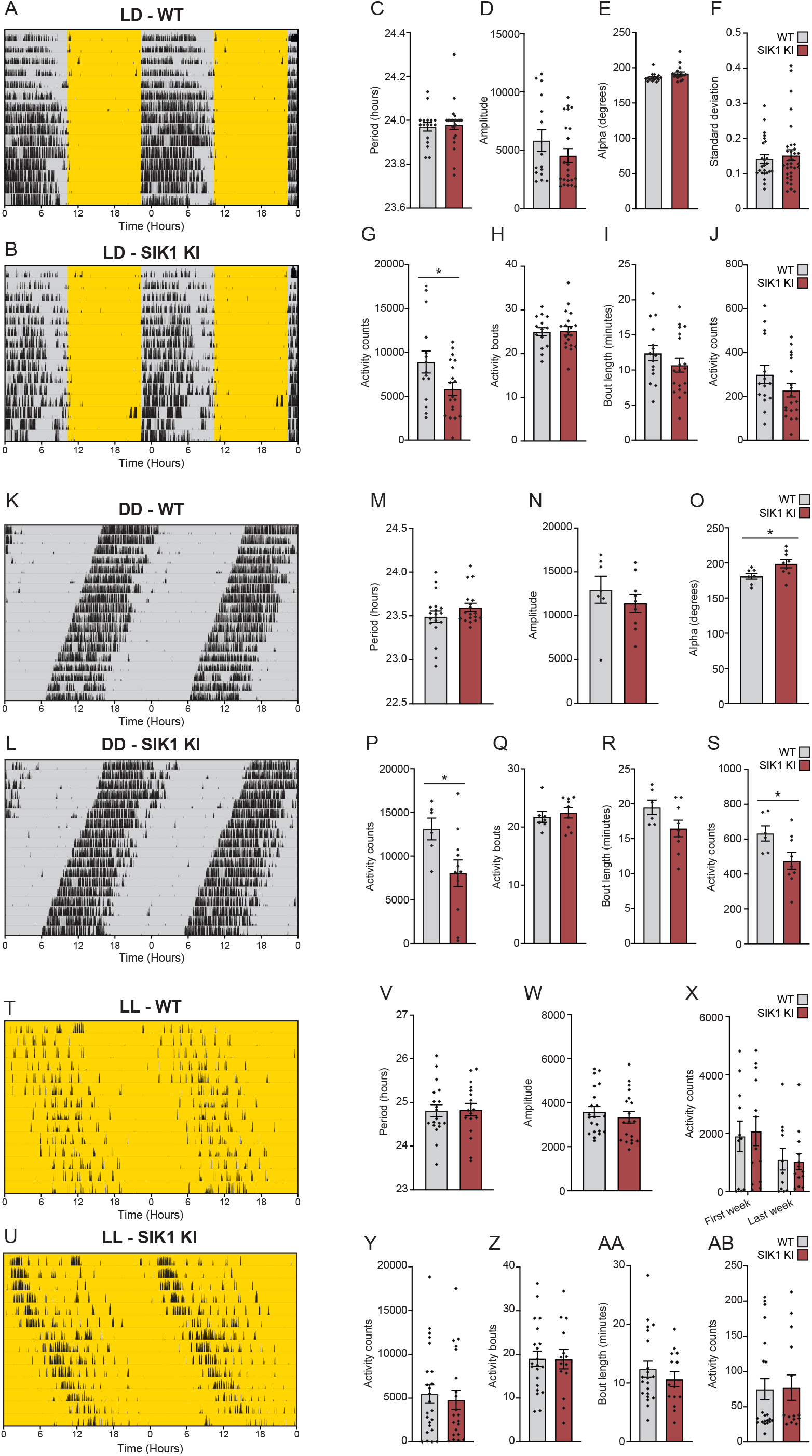
SIK1 KI animals have normal circadian activity and behaviour under a 12:12 light/dark cycle and under constant conditions. Actograms from (A) WT and (B) SIK1 KI mice under a 12:12 LD cycle. Lights on - yellow, lights off - grey, activity - black. Each horizontal line represents 24 hours double plotted. (C) LD circadian period, (D) periodicity amplitude, (E) active period length – alpha, (F) standard deviation of activity onset, (G) average total daily wheel revolutions, (H) average activity bouts per day, (I) average bout length and (J) average wheel revolutions per activity bout. Actograms from (K) WT and (L) SIK1 KI mice in DD. (M) DD circadian period, (N) periodicity amplitude, (O) active period length, (P) average total daily wheel revolutions, (Q) average activity bouts per day, (R) average bout length and (S) average wheel revolutions per activity bout. Actograms from (T) WT and (U) SIK1 KI mice housed under constant light (LL). (V) LL circadian period, (W) periodicity amplitude, (X) average total daily wheel revolutions in the first week under LL and the final week under LL, (Y) average total daily wheel revolutions over the entire LL period, (Z) average number of activity bouts per day, (AA) average bout length and (AB) average number of wheel revolutions per activity bout. Data are mean ± SEM, n = 13-33 (LD), 6-18 (DD) or 7-24 (LL). Statistical analysis was conducted by two-tailed student’s t-test. ns P > 0.05, * P < 0.05.

**Figure S2. Related to Figure 1.**
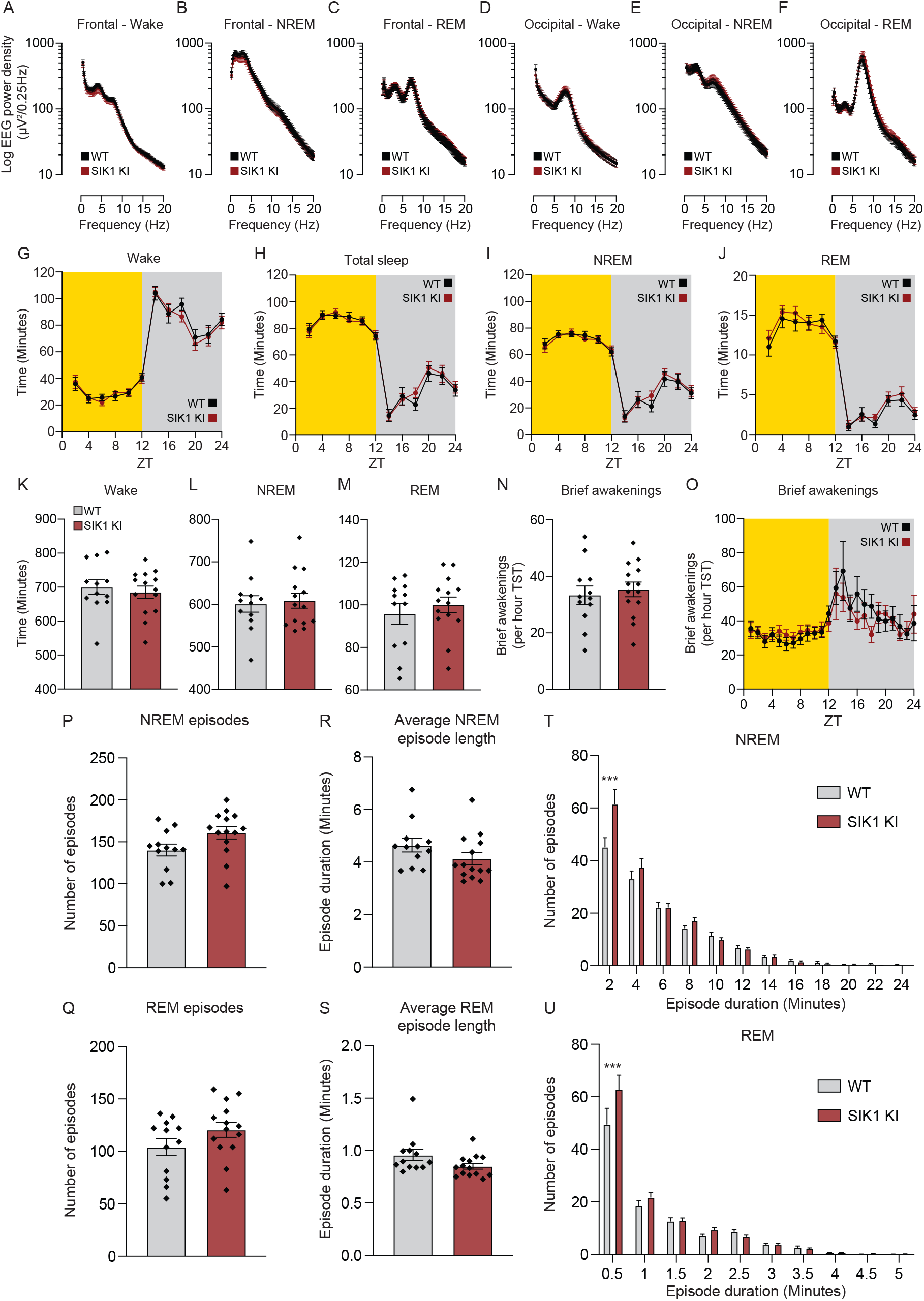
SIK1 KI animals have normal sleep architecture under stable entrainment. WT and SIK1 KI animals were housed under a 12:12 LD cycle with continuous sleep recording. Baseline average frontal EEG spectra during (A) wake, (B) NREM and (C) REM. Average occipital EEG spectra during (D) wake, (E) NREM and (F) REM. (G) Wake, (H) total sleep, (I) NREM and (J) REM in 2-hour bins, and (K) wake, (L) NREM and (M) REM over the entire baseline day. Brief awakenings per total hour sleep time (N) over the entire baseline day or (O) in 1-hour bins. Total (P) NREM and (Q) REM sleep episodes. Average (R) NREM and (S) REM episode duration. Histogram analysis of (T) NREM and (U) REM episode duration. Data are (A-S) Mean ± SEM or (T,U) mean + SEM. n = 12-14. Statistical analysis was conducted by (A-J, O, T, U) two-way ANOVA with Sidak’s correction or by (K-N, P-S) two-tailed student’s t-test. ns P > 0.05, *** P < 0.001.

**Figure S3. Related to Figure 1.**
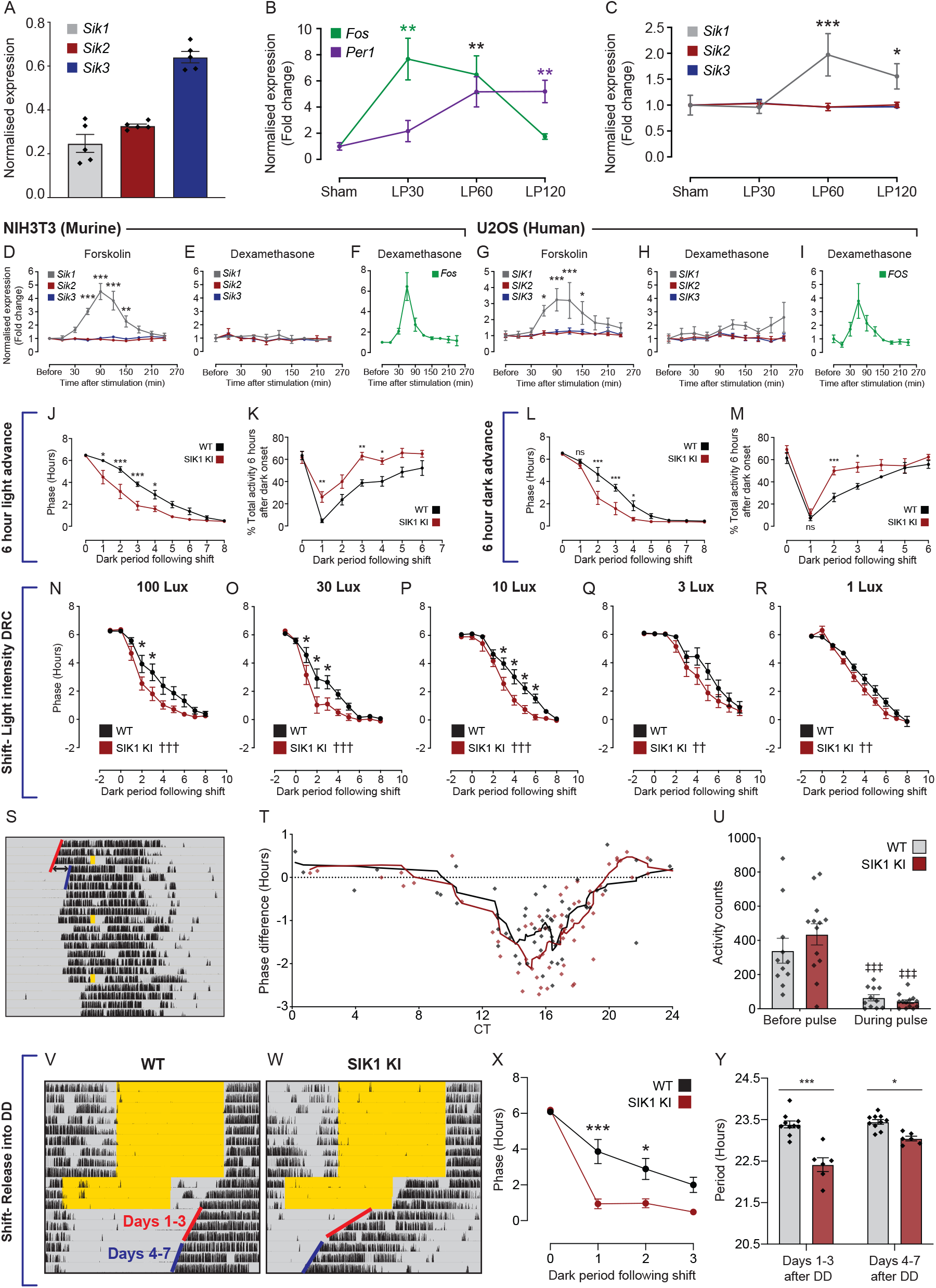
SIK1 is a light-inducible kinase that specifically regulates the speed of photic entrainment. (A) SCN tissue was collected at ZT16 and *Sik1, Sik2* and *Sik3* expression measured by RT-PCR. SCN expression of (B) *Fos* and *Per1*, and (C) *Sik1, Sik2* and *Sik3* in response to no light pulse (sham), a 30 min light pulse (LP30), and 30 and 90 min following the light pulse (LP60 and LP120, respectively). Expression of (D) *Sik1, Sik2* and *Sik3* in response to 10 μM forskolin, and expression of (E) *Sik1, Sik2, Sik3* and (F) *Fos* in response to 200 nM dexamethasone in murine NIH3T3 cells. Expression of (G) *SIK1, SIK2* and *SIK3* in response to 10 μM forskolin, and expression of (H) *SIK1, SIK2, SIK3*, and (I) *FOS* in response to 200 nM dexamethasone in human U2OS cells. (J) phase (activity onset minus dark onset) and (K) percentage of total activity within the first 6 hours after dark onset, following a 6-hour light advance of WT and SIK1 KI animals. (L) phase and (M) percentage of total activity, following a 6-hour dark advance. WT and SIK1 KI animals were subjected to a repeated LD shift paradigm where after each 6-hour LD cycle advance and subsequent re-entrainment, the light intensity was reduced by one third and the 6-hour advance repeated. Phase analysis of the (N) 100, (O) 30, (P) 10, (Q) 3 and (R) 1 lux periods. (W-Y) WT and SIK1 KI animals were housed in DD and subjected to multiple 30 min, 100 lux nocturnal light pulses (S, actogram with phase shift calculation highlighted). (T) Phase response curve analysis. (U) Average number of wheel revolutions in the 30 minutes before, and during, the light pulse. Actograms from (V) WT and (W) SIK1 KI animals, and (X) phase following a 6-hour light advance and release into DD on day 3. (Y) Circadian period of days 1-3 and 4-7, following release into DD. Data are mean ± SEM, n = (A-C) 5, (D-I) 3, (J-M) 6-10, (N-U) 9-13 and (V-Y) 6-10. Statistical analysis was conducted by (B-I) one-way ANOVA with Dunnett’s correction or (J-Y) two-way ANOVA with Sidak’s correction ns P > 0.05, * P < 0.05, ** P < 0.01, *** P < 0.001, either to control or comparing between the genotypes at the indicated time points. †† P < 0.01, ††† P < 0.001 for an overall genotype effect as determined by two-way ANOVA. ‡‡‡ P < 0.001 comparing activity counts before and after light pulsing.

**Figure S4. Related to Figure 2.**
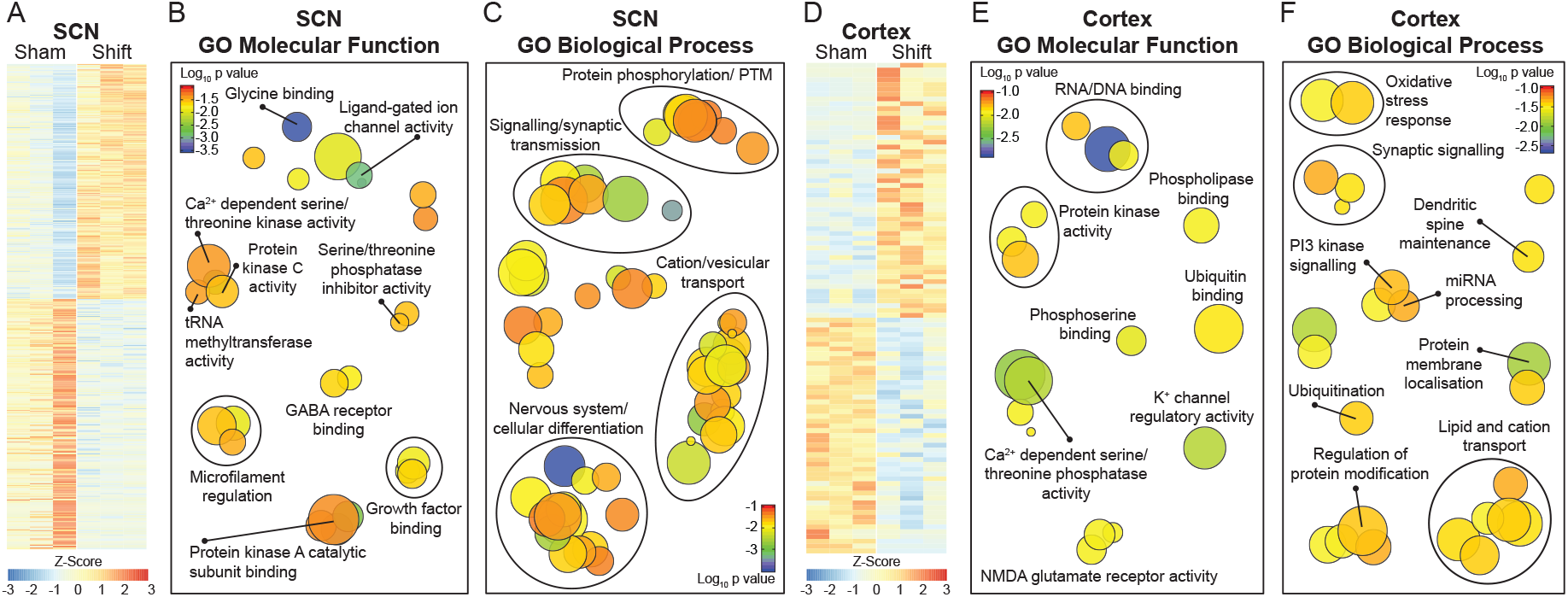
Nocturnal light exposure remodels the SCN and cortical transcriptome. SCN and cortex samples were collected either 6 hours into a 6-hour light advance (Shift) or at the same relative time, but without a light shift (Sham), and RNA sequencing performed. (A) Heatmap of differentially expressed SCN genes. (B) GO molecular function and (C) GO biological process pathway analysis of SCN differential genes. (D) Heatmap of differentially expressed cortex genes. (E) GO molecular function and (F) GO biological process pathway analysis of cortex differential genes. (A and D) Data are z score normalised per row. n = 3-6.

**Figure S5. Related to Figure 3 and Figure 4.**
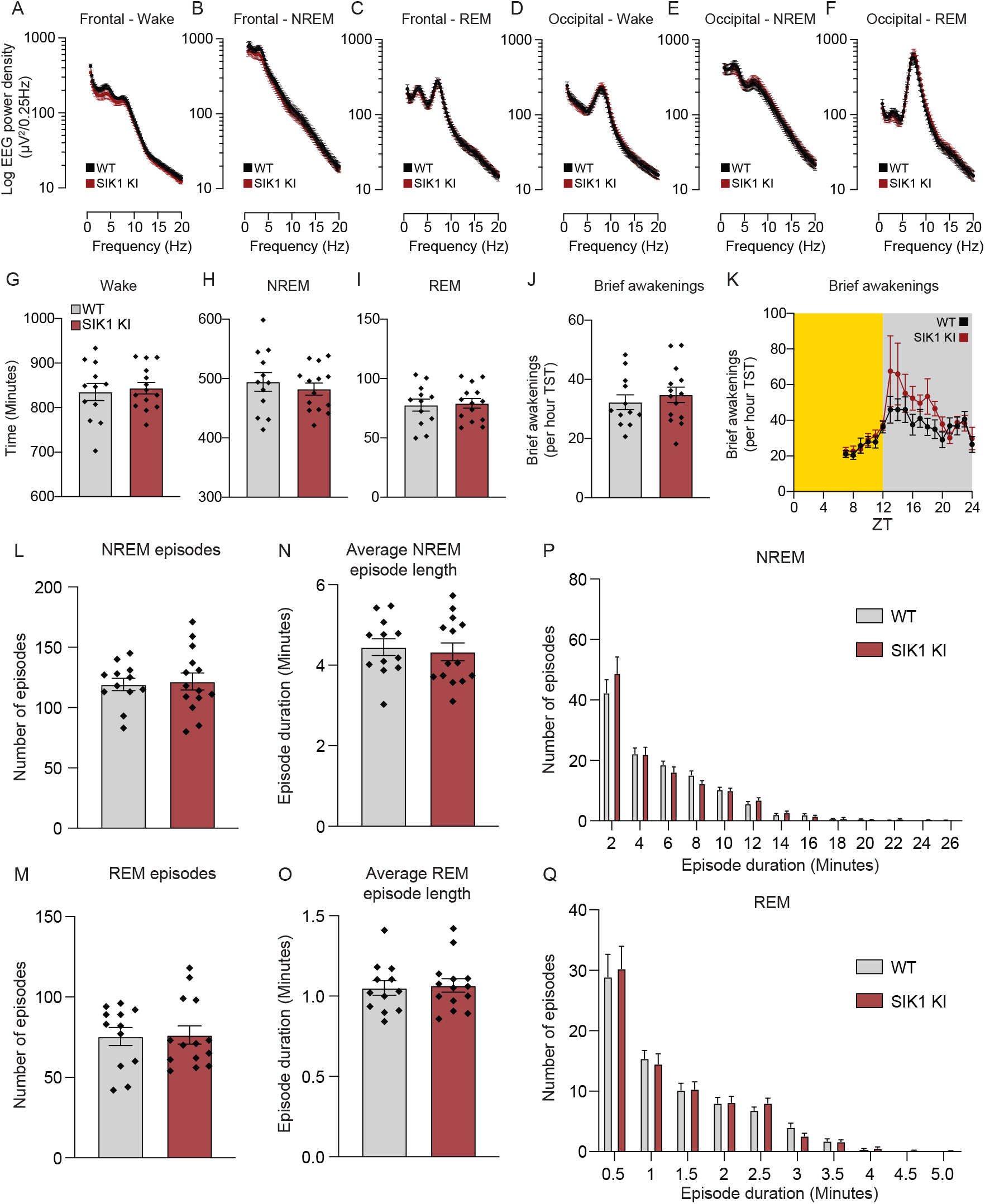
SIK1 KI animals have normal rebound sleep and sleep architecture following 6 hours of sleep deprivation. WT and SIK1 KI animals were housed under a 12:12 LD cycle with continuous sleep recording. Sleep deprivation was then conducted from ZT0 – ZT6. Average frontal EEG spectra during (A) wake, (B) NREM and (C) REM, and average occipital EEG spectra during (D) wake, (E) NREM and (F) REM over the entire sleep deprivation day. Total (G) wake, (H) NREM and (I) REM over the entire sleep deprivation day. Brief awakenings per total hour sleep time (J) over the entire sleep deprivation day or (K) in 1-hours bins. Total (L) NREM and (M) REM sleep episodes. Average (N) NREM and (O) REM episode duration. Histogram analysis of (P) NREM and (Q) REM episode duration. Data are (A-O) Mean ± SEM or (P,Q) Mean + SEM. n = 12-14. Statistical analysis was conducted by (A-F, K, P, Q) two-way ANOVA with Sidak’s correction or by (G-J, L-O) two-tailed student’s t-test. ns P > 0.05, *** P < 0.001.

**Table S1. RNA sequencing of sham and jet lag SCN and Cortex**

**Table S2. Whole brain quantitative phosphoproteomics of sham and LD cycle shifted WT and SIK1 KI mice**

**Table S3. Overlapping phosphoprotein lists from the LD cycle shift experiments**

**Table S4. Whole brain quantitative phosphoproteomics of sham and sleep deprived WT and SIK1 KI mice and phosphoprotein overlaps**

**Table S5. RT-PCR primer sequences**

## Notes

### Competing Interest Statement

The authors have declared no competing interest.

